# Genetic Loss of Nicotinamide Nucleotide Transhydrogenase Prevents from Cardiometabolic Heart Failure with Preserved Ejection Fraction

**DOI:** 10.1101/2023.04.21.537792

**Authors:** Mark E. Pepin, Sumra Nazir, Philipp J.M. Konrad, Friederike Schreiter, Matthias Dewenter, Johannes Backs

**Author notes:** **Correspondence:** Johannes Backs, MD, Institute of Experimental Cardiology, Medical Faculty Heidelberg, Heidelberg University, Im Neuenheimer Feld 669, 69120 Heidelberg, Germany. These authors contributed equally (Mark E. Pepin, Sumra Nazir, Philipp J.M. Konrad).

## Abstract

**Rationale:** Heart failure with preserved ejection fraction (HFpEF) represents a common clinical endpoint of cardiometabolic diseases which impair myocardial diastolic relaxation. Although myocardial redox perturbations are known to accompany HFpEF, the specific role of mitochondrial oxidative stress has not been demonstrated yet.

**Objective:** Based on an observation that C57BL6/N – but not C57BL6/J – mice develop diastolic dysfunction when provided an *ad libitum* high-fat and 0.5% N(ω)-nitro-L-arginine methyl ester (HFD+L-NAME) diet, we conducted a multi-cohort murine study to determine whether the loss of Nicotinamide Nucleotide Transhydrogenase (NNT), a mitochondrial transhydrogenase that couples NADPH:NADP^+^ to NADH:NAD^+^ homeostasis, protects mice from developing cardiometabolic alterations.

**Methods and Results:** Two cohorts of 12-week-old male and female mice possessing wild-type (*Nnt*^+/+^) or deleted (*Nnt*^-/-^) NNT were challenged by HFD+L-NAME for 9 weeks (n = 6-10). Male *Nnt*^+/+^ mice developed obesity (23.2% Δ, *P* = 0.003), arterial hypertension (24 ± 5 Δ mmHg, *P* = 0.023), impaired glucose tolerance (*P* = 0.006), and reduced maximal treadmill running distance (−172 ± 73.1 Δ m, *P* = 0.006) following 9 weeks HFD+L-NAME, whereas male *Nnt*^*-/-*^ mice did not. Female mice were protected from cardiometabolic dysfunction regardless of *Nnt* genotype. Cardiac functional and morphologic characterization revealed similar NNT-dependent and sex-specific increases in E/e’ (42.8 vs. 21.5, *P* < 0.001) and E/A (2.3 vs 1.4, *P* = 0.007) ratios, diastolic stiffness (0.09 vs 0.04 mmHg/μL, *P* = 0.02), and myocardial fibrosis (*P* = 0.02). Unsupervised transcriptomic analysis identified distinct genetic and dietary signatures, wherein *Nnt*^+/+^ exhibited disproportionate perturbations in various mitochondrial oxidative pathways following HFD+L-NAME. Our search for putative transcriptional regulators identified NNT-dependent suppression of NAD+ dependent deacetylase *Sirt3*.

**Conclusions:** Taken together, these observations support that the genetic disruption of *Nnt* protects against both cardiac and metabolic consequences of HFD+L-NAME, thus highlighting a novel etiology-specific avenue for HFpEF therapeutics.

## INTRODUCTION

Heart failure (HF) with preserved ejection fraction (HFpEF) is a high-prevalence clinical syndrome characterized by evidence of congestive heart failure attributable ventricular diastolic dysfunction and other cardiometabolic symptoms such as impaired exercise tolerance and insulin resistance. HFpEF is known for its poor clinical outcomes and lack of responsiveness to most guideline-directed HFrEF therapies.^1^ Consequently, HFpEF is widely considered a distinct clinical entity requiring its own therapeutic strategy.

Although the origins of HFpEF are multifactorial, mitochondrial oxidative stress is theorized to represent a common underlying pathogenesis. Epidemiologic studies have identified an array of genetic,^2,3^ lifestyle and environmental^4^ factors which ultimately lead to HFpEF as a clinical endpoint.^5–7^ They have been shown to disrupt mitochondrial homeostasis in HFpEF,^8^ ultimately yielding a metabolic signature that is distinct from HFrEF.^9^

Preclinical models of HFpEF seek to reproduce the individual – and synergistic – myocardial consequences of obesity,^10^ diabetes mellitus,^11^ hypertension,^12^ and aging.^13^ Among the models used to characterize HFpEF,^14^ the cardiometabolic features of HFpEF are most accurately mirrored by the combination of high-fat diet (HFD) feeding and N(ω)-nitro-L-arginine methyl ester (L-NAME), a chemical inhibitor of nitric oxide synthase.^15^ Despite its high-precision phenotyping, the molecular impact of this “two-hit” model (HFD+L-NAME) HFpEF model on mitochondrial oxidative stress signaling has not yet been shown.

The current multi-cohort study therefore explores whether NNT, an enzyme responsible for maintaining mitochondrial redox homeostasis, is required for HFD+L-NAME to induce both cardiometabolic and myocardial diastolic dysfunction. Taken together, our results reveal that intact *Nnt* expression is required for HFD+L-NAME to yield a cardiometabolic and myocardial HFpEF phenotype. Additionally, the current study underscores the importance of selecting an appropriate genetic sub-strain when designing a preclinical HFpEF study.

## METHODS

### Ethics

All animal experiments were validated using at least two independent cohorts (n = 3-5/cohort) and were performed in accordance with the European Community guiding principles in the care and use of animals (2010/63/UE, 22 september 2010) as authorized by the Regierungspräsidium Karlsruhe (G-42/21, Baden-Württemberg, Germany). Murine RNA-sequencing data have been deposited in NCBI’s Gene Expression Omnibus (GEO) and are accessible through GEO Series accession number GSE225557 (https://www.ncbi.nlm.nih.gov/geo/query/acc.cgi?acc=GSE225557).

### Animal Model

12-week-old male and female mice from C57BL/6N and C57BL/6J backgrounds (Janvier Labs) were bred at the Klinisch Experimenteller Bereich (KEB) mouse facility of Heidelberg University. All cohorts of mice were group-housed (≤4 mice/cage) on a 12:12-h light-dark cycle from 06:00 to 18:00 at 25 ± 1°C and constant humidity with *ad libitum* access to standard chow (2916, Teklad) or HFD (58.0 kcal% fat, D12492) and water. N(ω)-nitro-L-arginine methyl ester (L-NAME, 0.5 g/L, pH = 7.4, Sigma-Aldrich) was added to light-sensitive bottles of cages administered HFD, as previously described.^16^ This two-hit model is hereafter described as HFD+L-NAME throughout the manuscript. Prior to the experiment, C57BL/6J mice were backcrossed with C57BL/6N to eliminate the 5-exon *Nnt* deletion characteristic C57BL/6J mice. We detected by PCR of DNA extracted from tail clippings the exon-exon boundary lost in *Nnt*^-/-^ mice using the following primers: 5’-GTA GGG CCA ACT GTT TCT GC-3’ (fwd), 5’-TCC CCT CCC TTC CAT TTA GT-3’ (rvs, C57BL/6J), and 5’-CAG TCA ACA TTG CAG GTA G-3’ (rvs, C57BL/6N). Consequently, these genotypes are denoted *Nnt*^-/-^ and *Nnt*^+/+^ for substrains with impaired or intact *Nnt* expression, respectively.

### Glucose tolerance

7 weeks after initiating dietary treatment, glucose tolerance was assessed via intraperitoneal (i.p.) injection of glucose (2 g/kg, 20% wt/vol D-glucose [Sigma] in 0.9% wt/vol saline) in 4-hour fasted mice, where blood glucose was measured at 0, 15, 30, 60, and 120 minutes using a TheraSense Freestyle glucometer. To control for known circadian variation of glucose tolerance,^17^ fasting began at 6 AM CEST for all experimental cohorts.

### Exercise Tolerance

8 weeks after beginning the dietary regimen, endurance exercise capacity was assessed using a single bout of graded maximal treadmill running, as previously described.^18^ After a 3-day period of acclimatization to the treadmill, mice were placed on the treadmill at a 0° incline, and the shock grid was activated. Treadmill speeds were sequentially increased, starting at 9.6 m/min and by 1.8 m/s every 2 minutes thereafter. The treadmill test was terminated when mice sustained contact with the shock grid for 5 seconds, defined as the point of exhaustion.

### Cardiac Echocardiography and Pressure-Volume (PV) Loop Analysis

Non-invasive systolic cardiac parameters were estimated via 2-dimensional transthoracic echocardiography without anesthesia using a VisualSonics Vevo 2100 machine, with diastolic measurements taken with mice under sedation with isofluorane (1.0-1.5 vol %). Echocardiography parameters were estimated from tracings using the Fujifilm Visualsonics® Vevo LAB (v5.6.1) software.

### PV loop analysis and tissue isolation

Following 9 weeks of diet administration, mice were fasted for 2 hours prior to invasive cardiac functional characterization via PV loop analysis for 2 hours (starting at 0800). Trunk blood was collected, and hearts were harvested and snap-frozen at -80°C until analysis. Sections of heart and liver were fixed in ROTI®Histofix 4 % (cat. #5666.2) and parafilm-embedded for histological processing.

### Histology

Parafilm-embedded sections of ventricular and hepatic cross-sections were used for histologic analysis following PV loop analyses. Tissue was paraffin-embedded, cut, and stained with hematoxylin/eosin, Masson-Trichrome, and periodic acid Schiff (PAS) staining using standard techniques. Images were obtained using a light microscope at 20X magnification (Zeiss Axioskop 40, Oberkochen, Germany). Cardiomyocyte cross-sectional area was quantified via automated high-content screening using Axiovision LE software (Zeiss, Oberkochen, Germany).

### RNA-sequencing analysis

Isolated RNA from left ventricles of 21-week-old *Nnt*^+/+^ and *Nnt*^-/-^ mice was analyzed for quality using a Bioanalyzer 2100 (Agilent). Total RNA was depleted from ribosomal RNA, polyA-enriched, fragmented, and paired-end sequenced at the European Molecular Biology Laboratory (EMBL). We provide details of the computational tools and coding scripts used in the current study as an online supplement via GitHub data repository: https://github.com/mepepin/NNT_HFpEF. Alignment of reads were mapped to the latest *Ensembl* C57BL/6N annotation (c57bl_6NJ_107.gtf) to quantify both gene- and exon-level expression. Alignment was accomplished using *STAR* (2.7.10a), with differential gene expression calculated using *DESeq2* (1.32.0) within the R (4.1.2) statistical computing environment.^19,20^ In this way, we estimated dispersion via maximum-likelihood and minimized it using the empirical Bayesian method to provide normalized counts based on dispersion estimates and sample size. We then calculated differential expression from normalized read counts via Wald test with Bonferroni *post hoc* adjusted *P-*value for each aligned and annotated gene. Statistical significance was assigned via unpaired two-tailed Bonferroni-adjusted *P* < 0.05, with biological significance assumed at |Fold-Change| > 50% with normalized count sum > 1.

### Western Blot Analysis

To measure protein level alterations in response to high fat-diet and L-NAME feeding, snap-frozen left ventricular tissue was pulverized, and resuspended in RIPA buffer, incubated on ice for 30 minutes, and centrifuged at 4C for 10 minutes at 10,000g to remove insoluble material. Lysate was then diluted in Laemmli sample buffer and boiled for 5 minutes. 5–10 mg of sample was then subjected to SDS-PAGE on 10% acrylamide gels at a constant current equal to 20 mA per gel and transferred to a Protran nitrocellulose membrane (Whatman; Dassel, Germany) using the BioRad® semi-dry transfer apparatus at 100V for 1 hour. Membranes were blocked in 5% dry milk in TBST (Tris-buffered saline +0.1% Tween), and then incubated over night at 4C with appropriate primary antibody in TBST at 1:1,000. The membranes were then washed 3x in TBST before incubation for 1 hour at room temperature with peroxidase-conjugated secondary antibodies in TBST at 1:10,000 (Southern Biotech, #1031-05, #4050-05). Antibody binding was detected using an enhanced chemiluminescence HRP substrate detection kit (Millipore; Watford, UK). The following antibodies were used: anti-GAPDH (Chemocon, #MAB374), and anti-NNT (Abcam, #ab110352).

### Statistics

For all pairwise comparisons, the Shapiro-Wilk test for normality was performed to determine the most appropriate statistical test. All pairwise factors exhibiting a parametric distribution were evaluated using student t-test with Benjamini-Hochberg adjustment; otherwise, a Mann-Whitney test was used. All data are reported as mean ± standard deviation unless otherwise specified. Statistical analyses and data visualization were completed using GraphPad Prism version 9.3.1 for Macintosh (GraphPad Software, San Diego, CA) and R software, version 4.1.2 (R Foundation for Statistical Computing, Vienna, Austria). Statistical significance was assigned when *P* < 0.05 unless otherwise specified.

## RESULTS

### Strain-specific differences in cardiometabolic HFpEF Induction

The initial observation using commercial strains C57BL/6J and C57BL/6N was made using HFD+L-NAME feeding relative to control diet, which demonstrating robust differences in diastolic cardiac function (**Fig. 1A**). Among the genes known to differ between them,^21^ a 5-exon deletion was detected in the *Nnt* gene in the C57BL/6J strain. To determine whether disrupted *Nnt* expression is sufficient to prevent HFD+L-NAME associated HFpEF, we back-crossed C57BL/6J into the C57BL/6N substrain, and we and generated two mouse strains, *Nnt*^+/+^ and *Nnt*^-/-^ based on genotyping for the *Nnt* deletion (**Fig. 1B**). Quantification of myocardial RNA validated the presence of exon 7-11 (**Fig. 1C**), seen at the level of gene-wide RNA expression (*P* = 9.8 × 10^−52^) (**Fig. 1D**) and NNT full-length protein (**Fig. 1E**) in *Nnt*^+/+^ but not *Nnt*^-/-^.

**Figure 1.**
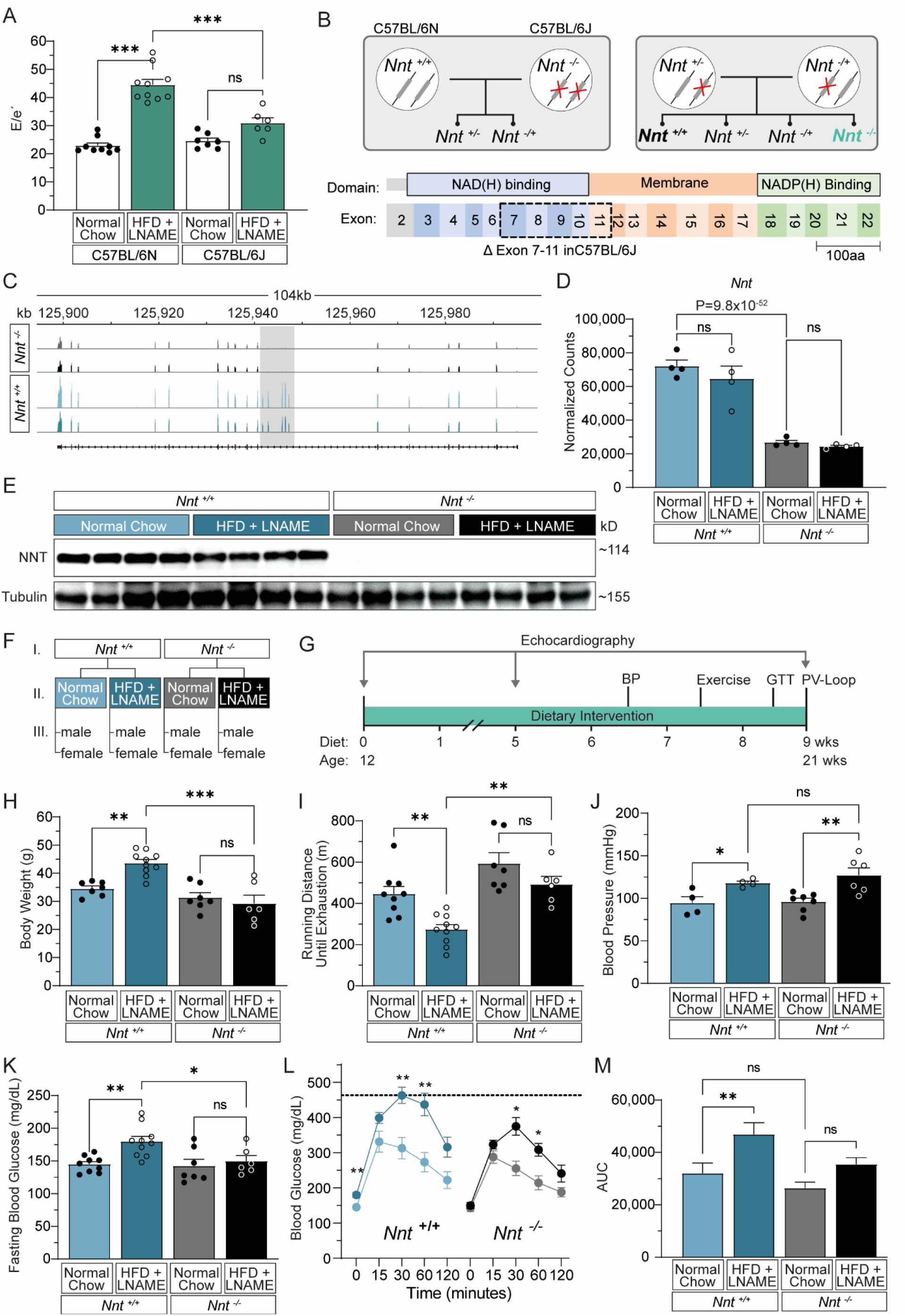
Genetic background unmasks an *Nnt*-dependent cardiometabolic response to HFD+L-NAME. **(A)** Non-invasive echocardiographic quantification of pulse-wave velocity to evaluate E/e’ in C57BL/6N and C57BL/6N mice following initiation of HFD+L-NAME vs normal chow diets (n = 10). **(B)** Illustration of *Nnt* mutation and breeding strategy to create *Nnt*^+/+^ mice not harboring the 5-exon *Nnt* deletion seen in C57BL/6J mice (*Nnt*^-/-^). **(C)** *Nnt* expression from bulk-tissue RNA-sequencing of left ventricle tissue (n = 4). Schema illustrating the experimental groups used in the study. **(D)** Integrated Genome Browser (IGV) depicting group averaged exon-level abundance of reads annotating to *Nnt* gene locus, highlighting absence of reads aligning to exons 7-11 among *Nnt*^-/-^ samples. **(E)** Western blot of NNT with corresponding reference protein Tubulin (n = 4). **(F)** Schema illustrating the experimental groups used in the study. **(G)** Experimental timeline with concordant endpoints used. **(H)** Body weight (grams) measured after 9 weeks of HFD+L-NAME or standard chow (CON) feeding in male mice. **(I)** Exercise capacity as assessed by graded treadmill running, recording total distance travelled (meters) until exhaustion. **(J)** Non-invasive systolic blood pressure (mm Hg) measured via tail-cuff at 6 weeks of HFD+L-NAME treatment. **(K)** Circulating glucose levels (mg/dL) measured via POC glucometer following 4-hr fasting. **(L)** Circulating glucose levels (mg/dL) during glucose tolerance testing of *Nnt*^+/+^ and *Nnt*^-/-^ mice.* **(M)** Area-under-curve analysis of glucose tolerance. Statistical significance was assigned via 1-way ANOVA with Tukey’s multiple comparisons test for all pairwise analysis, reporting group mean ± S.E.M. * P < 0.05, ** P < 0.01, *** P < 0.001

### Cardiometabolic dysfunction in HFD+L-NAME diet

To test whether the “two-hit” HFpEF model causes NNT-dependent cardiometabolic phenotype, *Nnt*^+/+^ versus *Nnt*^-/-^ mice (n = 6-10) were given *ad libitum* access to HFD and L-NAME (HFD+L-NAME) or standard chow (CON) for a total of 9 weeks beginning at 12 weeks of age (**Fig. 1F**), after which they underwent testing to quantify cardiac and metabolic characteristics (**Fig. 1G**). As expected, body weight increased by 18.2% among male *Nnt*^+/+^ mice after 5 weeks (*P* = 0.04) of HFD+L-NAME feeding, which further increased to 23.2% (*P* = 0.003) at 9 weeks (**Fig. 1H**). By contrast, male *Nnt*^-/-^ mice exhibited no weight gains over this 9-week period. Treadmill running endurance showed a similar trend, wherein male *Nnt*^+/+^ mice exhibited lower endurance exercise capacity (**Fig. 1I**). Although non-invasive blood pressure measurement via tail cuff demonstrated increased systolic pressure in male mice of both backgrounds (**Fig. 1J**), 4-hr fasting blood glucose measurements revealed diabetic-range glucose in male *Nnt*^+/+^ mice, whereas no differences were seen in *Nnt*^-/-^ mice (**Fig. 1K**). Similarly, glucose tolerance revealed a higher peak glucose concentration relative to *Nnt*^-/-^ males (**Fig. 1L**), as well as significantly increased glycemic area-under-curve (AUC) only in male *Nnt*^-/-^ relative to CON (*P* = 0.006, **Fig. 1M**). As has been published previously, female mice exhibited no alterations in weight (**Fig. S1A**), exercise tolerance (**Fig. S1B**), systolic blood pressure (**Fig. S1C)**, or glucose tolerance following 9 weeks of HFD+L-NAME regardless of genotype (**Fig. S1D-F**). Taken together, these data suggest both a sex-specific and NNT-dependent role of HFD+L-NAME on cardiometabolic dysfunction observed in this HFpEF model.

### Cardiac phenotyping reveals NNT-dependent diastolic dysfunction in HFD+L-NAME

To non-invasively validate the effect of HFD+L-NAME on cardiac dysfunction, we used systolic and diastolic echocardiography prior to initiating dietary intervention, at the 5-week, and at 9-week timepoints (**Fig. 2A**). Most notably, HFD+L-NAME produced higher atrial filling pressure by 5 weeks, where only *Nnt*^+/+^ mice displayed markedly higher ratios of early mitral inflow and mitral annular early diastolic velocities (E/e’, **Fig. 2B**, *P* < 0.001) and peak velocities from early-to-late diastole (E/A, **Fig. 2C**, *P* = 0.007), as estimated via pulse-wave and tissue doppler echocardiography, respectively. Similarly, pressure-volume (PV) loop analysis at 9-week harvest exposed a greater slope of the end-diastolic pressure-volume relationship (EDPVR, **Fig. 2D**), depicting changes known to occur in HFpEF only in *Nnt*^+/+^ – not *Nnt*^-/-^ – male mice (**Fig. 2E-F**).^22^ No evidence of diastolic dysfunction was found in female mice (**Fig. S2**). Two-dimensional M-mode echocardiography revealed no differences were seen in fractional shortening (**Fig. 2G**), although LV posterior (**Fig. 2H**) and anterior (**Fig. 2I**) wall thickness increased in male *Nnt*^+/+-^mice following 5 and 9 weeks HFD+L-NAME treatment; by contrast, ventricular dimensions and function were unchanged in *Nnt*^-/-^ male mice at any timepoint. Taken together, these data support an NNT-dependent and sex-specific influence of HFD+L-NAME on left ventricular diastolic dysfunction consistent with a compensated stage of the HFpEF syndrome.

**Figure 2.**
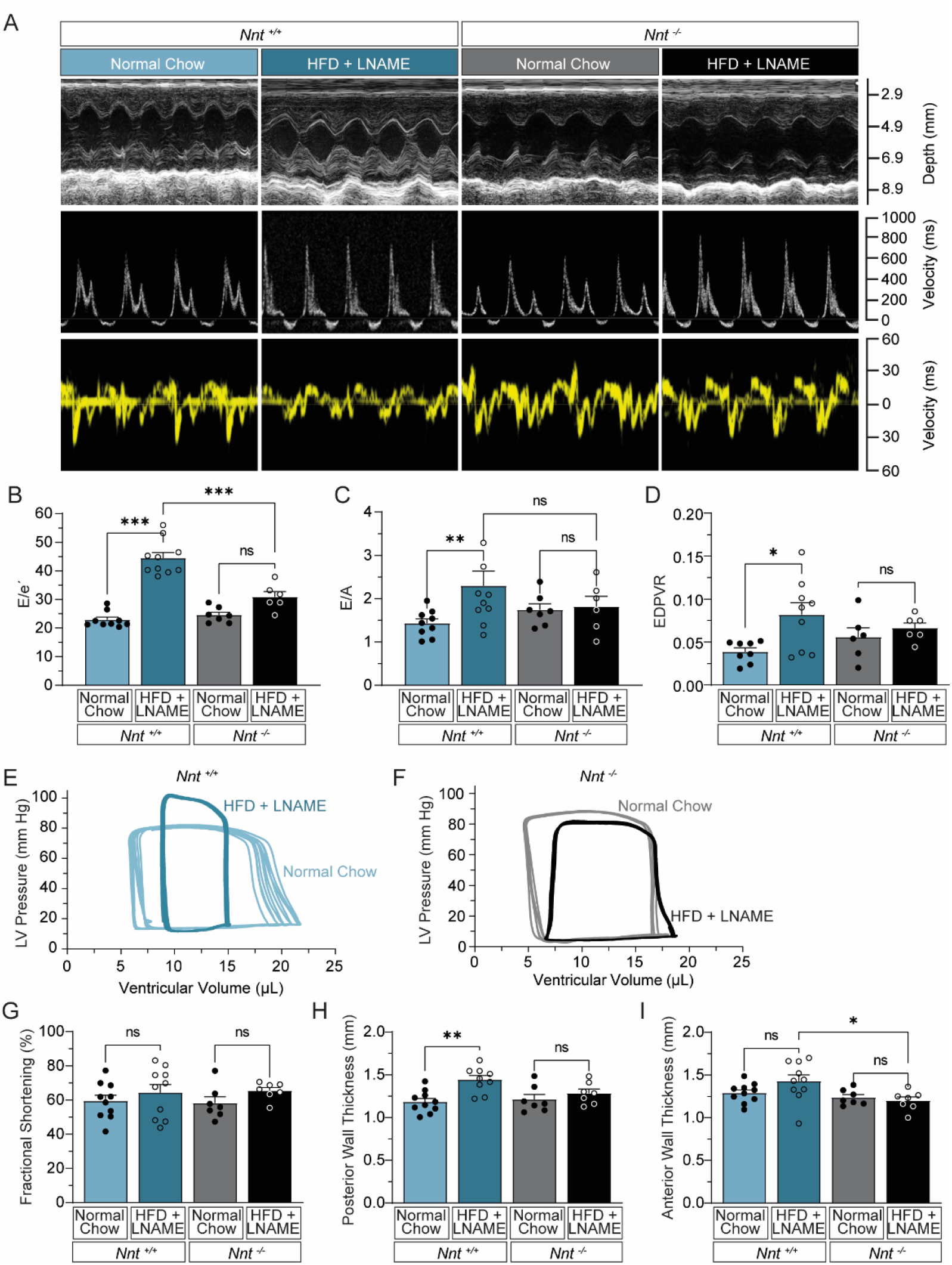
Cardiac phenotyping of HFpEF following HFD+L-NAME treatment. **(A)** representative images of non-invasive cardiac imaging via echocardiographic M-mode, pulse-wave, and tissue doppler studies for each group (n = 6-10). Quantified diastolic parameters of **(B)** E/e’ ratio (pulse-wave), **(C)** E/A ratio using pulse-wave and tissue doppler, and **(D)** slope of end-diastolic pressure-volume relationship using estimated exponent (Pressure = α x eβ x Volume + exponent) method. **(E - F)** Representative illustration of pressure-volume relationship in response to HFD+L-NAME diet relative to CON in *Nnt*^+/+^ and *Nnt*^-/-^ mice, respectively. **(G)** M-mode systolic parameters for left ventricular fractional shortening (%), **(H)** posterior wall thickness (mm), and **(I)** Anterior wall thickness (mm). Statistical significance was assigned via 1-way ANOVA with Tukey’s multiple comparisons test for all pairwise analysis, reporting group mean ± S.E.M. * P < 0.05, ** P < 0.01, *** P < 0.001.

### Morphologic evidence of myocardial hypertrophy and fibrosis in HFD+L-NAME

Histologic sectioning was performed to quantify morphologic differences in response to HFD+L-NAME. Although we found no differences in heart size (**Fig. 3A**) or biventricular weight (**Fig. 3B**) across the experimental groups, hearts of *Nnt*^+/+^ mice underwent significant increases in left ventricular fibrosis following HFD+L-NAME, which was not seen in *Nnt*^-/-^ mice (**Fig. 3C-D**). Although we saw no evidence of pulmonary edema, as was noted in the original report by Schiattarella *et al*.,^16^ exercise intolerance is considered a more sensitive – and clinically relevant – proxy of subclinical pulmonary congestion in compensated HFpEF.^23^

**Figure 3.**
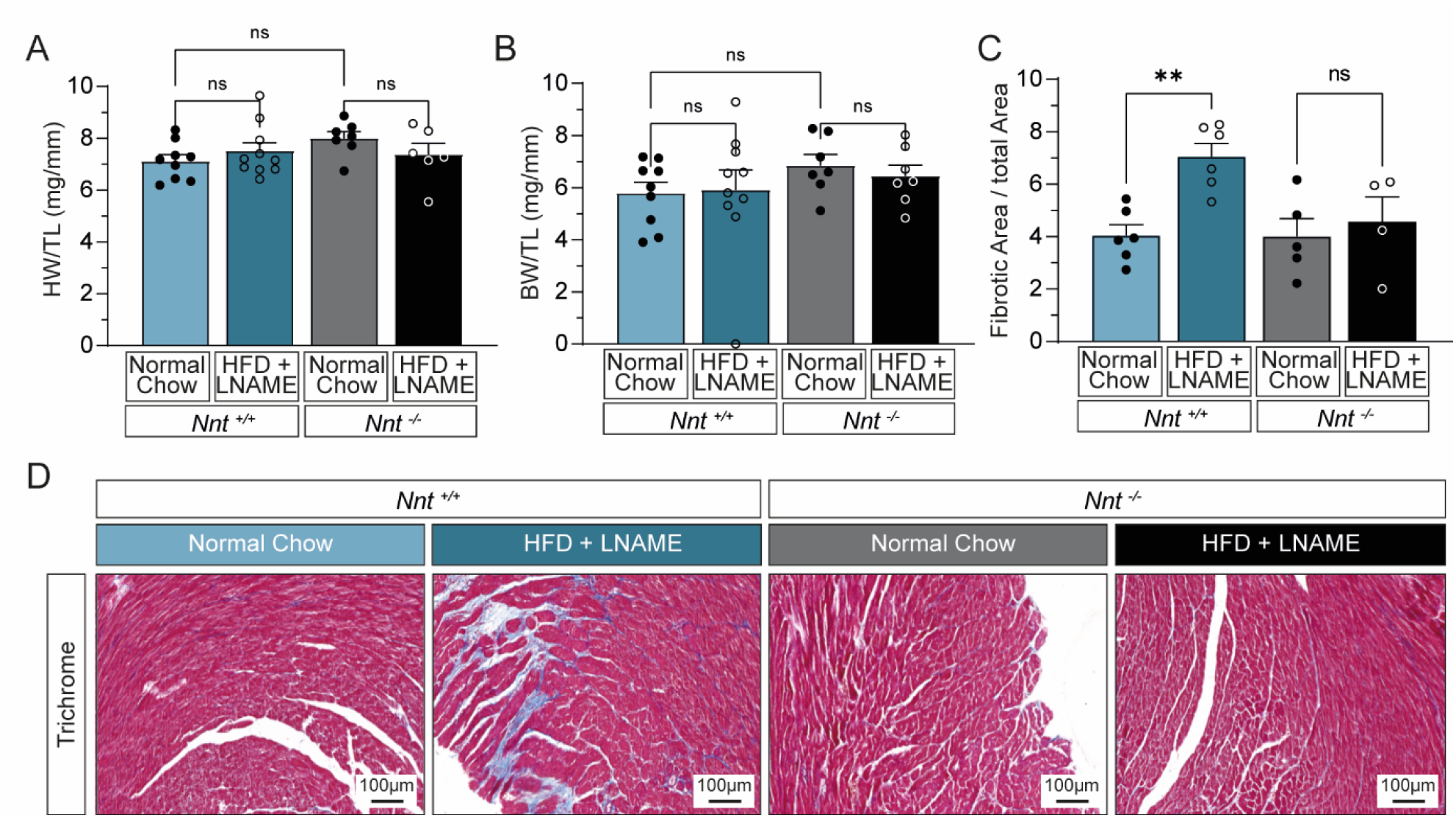
Loss of NNT attenuates perivascular fibrosis and oxidative stress leading to HFpEF following HFD+L-NAME. **(A)** Representative histologic images and **(B)** colorimetric quantification of left ventricles stained with trichrome from male *Nnt*^+/+^ and *Nnt*^-/-^ treated with normal chow or HFD + L-NAME (n = 6). **(C)** Heart weight normalized to tibia length (mg/μL). **(D)** Biventricular weight normalized to tibia length (mg/μL). Statistical significance was assigned via 1-way ANOVA with Tukey’s multiple comparisons test for all pairwise analysis, reporting group mean ± S.E.M. * P < 0.05, ** P < 0.01, *** P < 0.001.

### NNT-dependent myocardial gene expression

To identify the effects of HFD+L-L-NAME on myocardial gene expression, RNA-sequencing analysis of left ventricular tissue was performed on all experimental groups. We used a multidimensional scaling as an unsupervised clustering method, which uncovered robust signature of genotype-specific (PC1) and dietary (PC3) effects (**Fig. 4A**). Differential gene expression identified 1,798 differentially-expressed genes (DEGs) and 2,129 DEGs following 9 weeks of HFD+L-NAME among *Nnt*^+/+^ and *Nnt*^-/-^, respectively (*P* < 0.05). Among these, 229 were both increased and 124 decreased in both genotypes (**Fig. 4B**). Two-way analysis of differential expression via genotype and diet identified distinct patterns of gene expression, illustrated by hierarchical clustering and heatmap visualization, from which 459 *Nnt*-dependent and HFD+L-NAME responsive DEGs were identified (**Fig. 4C**).

**Figure 4.**
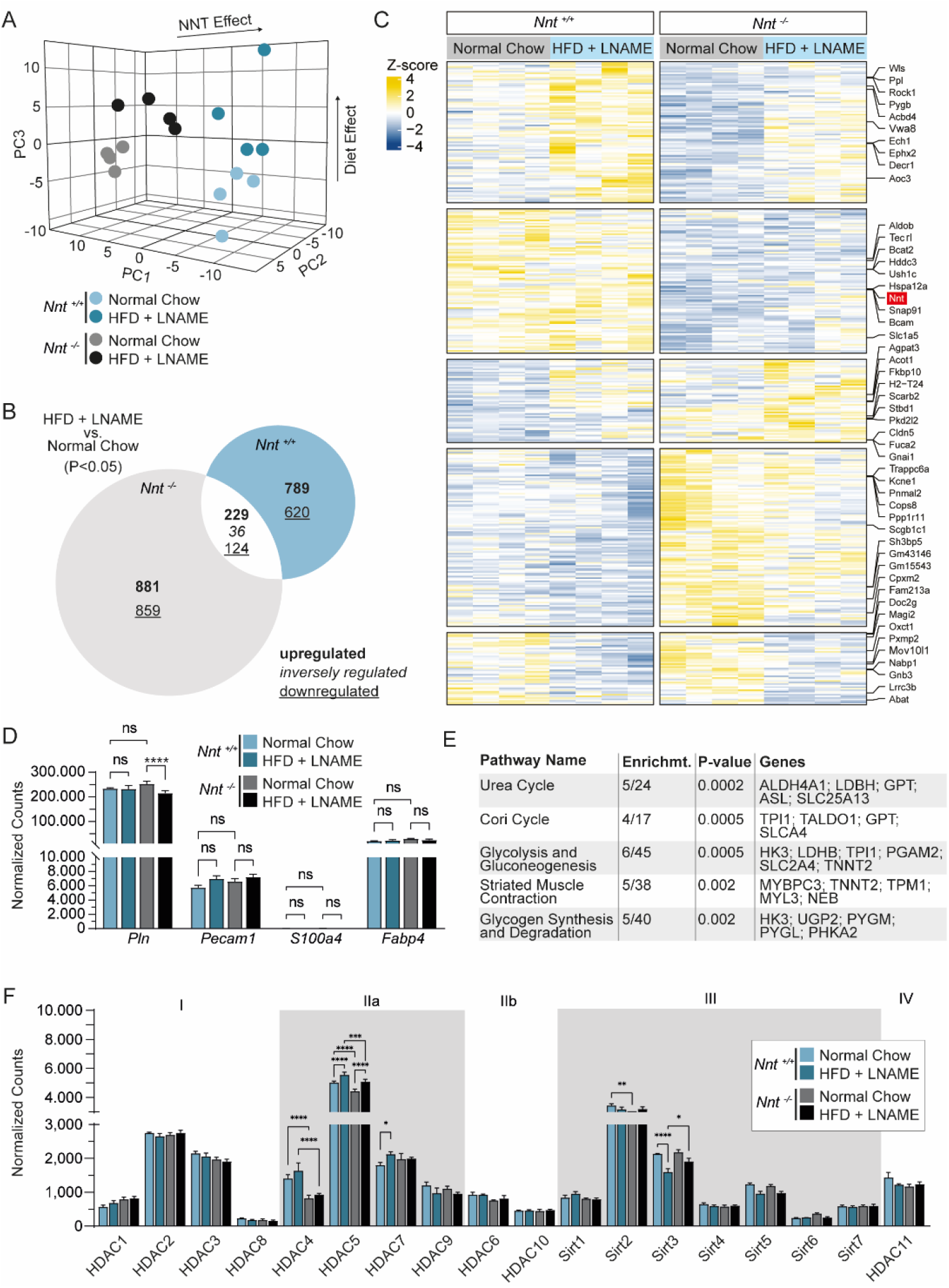
Distinct transcriptomic signature of HFD+L-NAME and NNT genotype. **(A)** 3-dimensional plot illustrating the multidimensional scaling (MDS) of top 10,000 genes based on RNA-sequencing of *Nnt*^-/-^ vs *Nnt*^+/+^ given normal chow (CON) or HFD+L-NAME (n = 4). The two principal components that account from the largest variance in DNA methylation were used to generate a scatterplot, flanked by density plots of each principal component. **(B)** Venn-diagram illustrating distinct and overlapping DEGs within *Nnt*^+/+^ and *Nnt*^-/-^ genotypes in after 9 weeks HFD+L-NAME feeding relative to CON (*P* < 0.05, | Fold-Change| > 1.5). **(C)** Heatmap and hierarchical clustering of differentially-expressed genes (DEGs) based on LRT analysis of Diet and Genotype (P < 0.05). **(D)** Gene expression of cell markers from RNA-sequencing (n = 4). **(E)** Gene-set enrichment analysis of *Nnt*-dependent gene expression in HFD+L-NAME relative to CON diet. **(F)** Gene expression of Class III histone deacetylases (Sirtuins), demonstrating exclusive dysregulation of *Sirt3*. **(G)** Gene expression of class I and class II histone deacetylases (HDACs). Statistical significance was assigned via 1-way ANOVA with Tukey’s multiple comparisons test for all pairwise analysis, reporting group mean ± S.E.M. * P < 0.05, ** P < 0.01, *** P < 0.001.

Variable tissue composition has been shown to influence differential expression analysis.^24,25^ Therefore, to assess the potential for morphological impact on the myocardial transcriptome, we used cardiomyocyte-specific (TNNT2, ACTN2, PLN), fibroblast-specific (TSLP, S100A4, PDGFRA), endothelial cell-specific (PECAM1, DACH1, ICAM2), and adipocyte-specific (HOXC8, HOXC9, FABP4) gene markers (**Figure 4D**).^26–28^ This exposed a high-degree of cardiomyocyte gene marker expression relative to other cell markers, and no dietary differences were noted. Owing to the importance immune cell composition on cardiac phenotype,^29,30^ *in-silico* method of cellular deconvolution was performed using *Xcell*^25^ to infer relative enrichment of immunologic and somatic cell-types in our sample set (**Supplemental Figure S3**); again, no group-wise differences were detected to confound our analysis. Gene set enrichment analysis of NNT-dependent HFD+L-NAME DEGs identified metabolic and mitochondrial pathways, including the Urea cycle (*P* = 0.0002), Cori Cycle (*P* = 0.0005), Glycolysis and Gluconeogenesis (*P* = 0.0005), Striated Muscle Contraction (*P* = 0.002), and Glycogen Synthesis and Degradation (*P* = 0.002) (**Fig. 4E**).

Because non-genomic activity of histone deacetylases (HDACs) have been implicated in the pathogenesis of HFpEF,^31^ we examined all Class I-III HDACs to determine whether our data reflected NNT dependence and/or cardiometabolic responsiveness to the HFD+L-NAME model. We identified *Sirt3* to be suppressed (1.34-fold, *P* = 0.0001) following HFD+L-NAME treatment in an NNT-dependent manner as the only histone deacetylase to exhibit this pattern across classes (**Fig. 4F**). However, we also found *Hdac5* as induced (P = 0.0) by HFD+L-NAME in both *Nnt*^+/+^ and *Nnt*^-/-^, whereas *Hdac4* was suppressed in *Nnt*^-/-^ (−1.4-fold, *P* = 0.02) regardless of diet.

## DISCUSSION

It is well established that murine sub-strains profoundly impact the recapitulation of human physiologic – and pathologic – responses to stress, and C57BL/6J and C57BL/6N are no exception; they have been bred separately for over 220 generations, amassing SNPs and indels that now punctuate their respective genomes.^32^ Consequently, they encompass a range of cardiovascular,^33–35^ metabolic,^36,37^ and even behavioral/neurologic phenotypes.^38,39^ The C57BL/6J sub-strain harbors a five-exon (7 – 11) deletion within *Nnt*, unlike the intact gene found among those from Taconic, Jackson Lab, Charles River or Janvier (C57BL/6N), a gene encoding the nicotinamide nucleotide transhydrogenase (NNT). As a key mitochondrial enzyme involved in NADPH production,^40,41^ the current study supports our hypothesis that metabolic stress converges onto mitochondrial ROS homeostasis to produce cardiometabolic HFpEF.

The 5-exon deletion of *Nnt* in C57BL/6J mice results in loss of function of the inter-conversion of NADPH and NADH.^42^ NNT couples mitochondrial NADH reservoir supplanted by oxidative phosphorylation to replenish nearly 50% of the mitochondrion’s NAPDH that is oxidized under aerobic conditions, thus maintaining a redox homeostasis that spans the heart’s physiologic range of metabolic operating conditions.^43,44^ Although this inter-conversion typically favors replenishment of NADPH, certain pathophysiological conditions of myocardial metabolic and/or oxidative stress reverse these dynamics to catalyze the reduction of NAD+ by de-protonating NADPH and – consequently – predisposing the heart to a pro-oxidative state.^45^ The current study therefore offers seminal evidence that pathogenesis of nitrosative stress-associated HFpEF requires intact NNT to reproduce the phenotypic features of HFpEF. Similarly, Schiattarella *et al*. used C57Bl/6N mice for the initial establishment of this HFD+L-NAME model.^16^ Intriguingly, although this initial report has been meticulously reproduced by several groups,^46,47^ other unpublished reports were unable to reproduce it, prompting the initial inquiry of whether genetic sub-strain selection accounts for this phenotypic variation.

### ROS and oxidative stress in HFpEF

Our study supports the emerging theory that cardiometabolic HFpEF is fundamentally a mitochondrial disorder that culminates from perturbations in ROS homeostasis.^48,49^ One such prior study has shown that myocardial cyclic guanosine 3’,5’-monophosphate (cGMP) concentrations are lower in HFpEF patients, and are inversely correlated with nitrotyrosine,^50^ which is a byproduct of nitrosative stress.^51^ Conversely, SGLT2 inhibitors lower myocardial oxidative stress in HFpEF,^52^ which may explain their clinical benefit.^53^

### Deacetylase-dependent basis of cardiometabolic HFpEF

Our analysis identified *Sirt3* as myocardial gene suppressed by HFD+L-NAME in an NNT-dependent fashion. Deletion of SIRT3 is known to cause hyperacetylation – and consequently, increased enzymatic activity – of mitochondrial metabolic intermediate enzymes.^54^ Over 60% of mitochondrial proteins contain acetylation sites,^55^ most of which regulate mitochondrial bioenergetics, and hyperacetylation of these enzymes has been linked to the HFpEF phenotype in mice.^56^ Specifically, nongenomic Class III HDACs SIRT1 and SIRT3 regulate Redox Homeostasis during Ischemia/Reperfusion in the Aging Heart.^57^ In support of this hypothesis, the Hill lab reported a causal involvement of *Sirt3* in cardiomyocytes for the development of HFpEF.^47^ Moreover, the differential expression patterns of some members of other deacetylases (HDACs) may also support a role and therapeutic potential of those HDACs in the pathogenesis of HFpEF.^58^

### Limitations

Although HFpEF is becoming an increasingly-prevalent condition in women, few – if any – murine models of HFpEF simulate this predisposition.^59^ One exception to this is a model developed by the de Boer laboratory,^60^ which used antiogensin infusion + high-fat feeding in 2 year old aged mice. Our data replicate the observation that 12 week old female mice are protected against diastolic dysfunction in response to HFD+L-NAME^61^; however, female gender portends a higher risk for HFpEF in humans.^62^ Although little is known regarding the sex-based cardiometabolic responses to high-fat feeding, it is feasible – yet untested – that hormonal factors confer cardioprotection to premenopausal females, particularly since the elevated HFpEF risk is only noted among postmenopausal women.^63^ Nevertheless, our data provide reproducible evidence that the “two-hit” murine HFpEF model is unsuitable for sex-based studies of human HFpEF regardless of genetic substrain.

## CONCLUSION

Our multicohort study reproducibly establishes sex-specific and NNT*-*dependent cardiometabolic manifestations of the “two-hit” HFpEF model. Beyond its insights into the utility of this pre-clinical HFpEF model, we aim to refocus preclinical models onto the mapping of precise clinical conditions whereby HFpEF develops, thus permitting the study of underlying pathophysiology that manifests in its end-stage as cardiac diastolic dysfunction leading to heart failure. Future studies should therefore attempt to stratify HFpEF into etiology-specific pathogenesis of myocardial dysfunction, which will yield more clinically effective approaches to the diagnosis, prevention, and treatment of human HFpEF.

## Acronyms

L-NAME: N(ω)-nitro-L-arginine methyl ester;
HFpEF: Heart failure with preserved ejection fraction;
NNT: nicotinamide nucleotide transhydrogenase;
HFD: high-fat diet;
CON: standard chow diet

## ACKNOWLEDGMENTS

We thank Joshua Hartmann, Sabine Kuß, Ina Broll, and Jutta Krebs for their technical scientific support and Marco Hagenmüller for managing the animal protocols.

## DATA AVAILABILITY

All data and analyses will be made available upon reasonable request to the corresponding author. RNA-sequencing sequencing data have been deposited in NCBI’s Gene Expression Omnibus (GEO), and are accessible through GEO Series accession number GSE225557 (https://www.ncbi.nlm.nih.gov/geo/query/acc.cgi?acc=GSE225557).

## CONTRIBUTIONS

M.E.P. designed and supervised the experiments, performed analysis, and wrote the manuscript. S.N. and M.E.P. performed echocardiography, harvested tissue, and managed mouse husbandry. P.J.M.K. analyzed performed molecular processing and histology. J.B. and M.E.P. conceived the current project and provided oversight regarding funding, interpretation, and revision of the manuscript. All authors read, edited, and approved the final manuscript.

## FIGURE LEGENDS

**Supplemental Figure 1.**
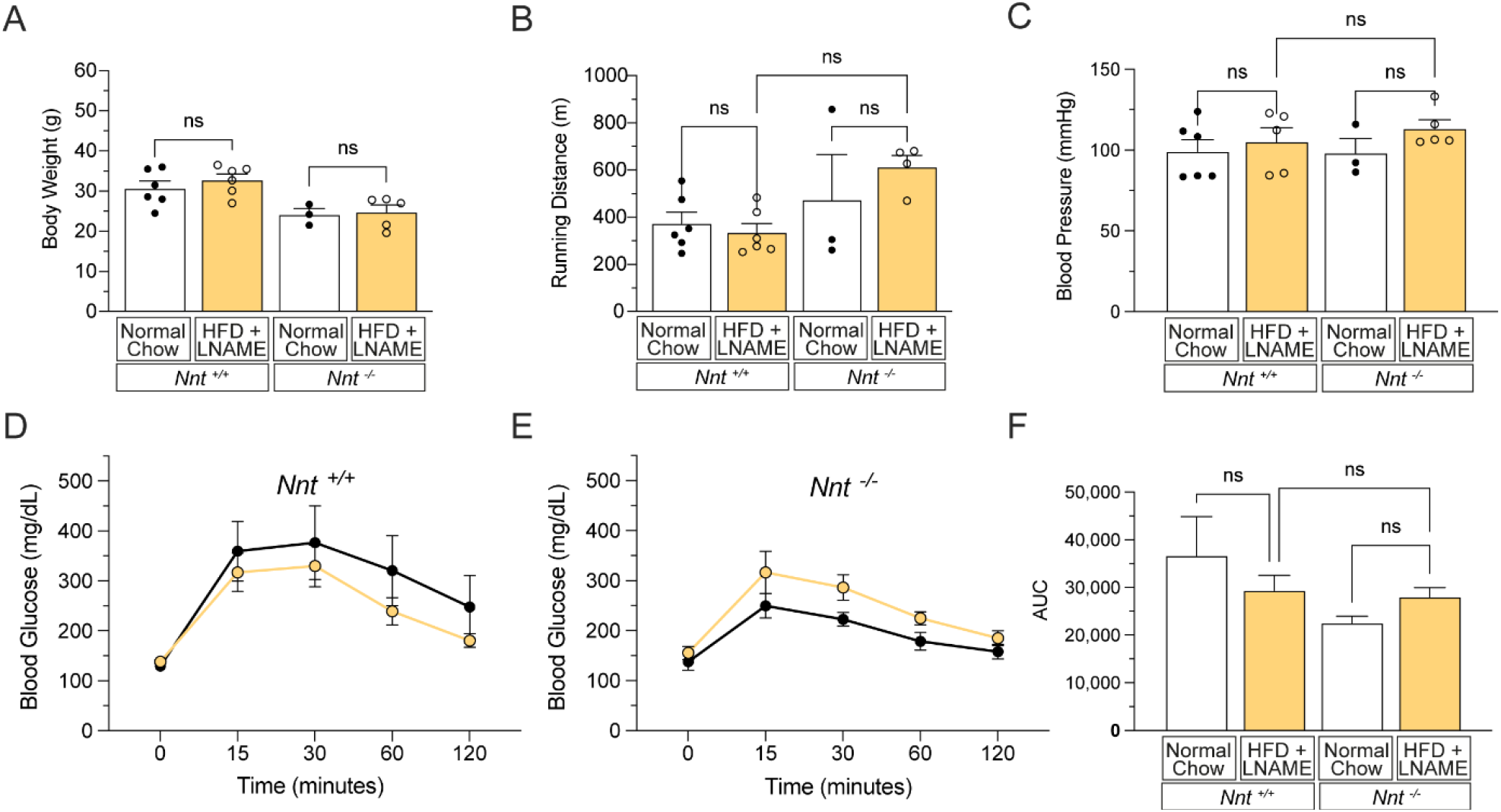
Lack of metabolic phenotype in female mice independent of *Nnt* genotype. **(A)** Body weight (grams) measured after 9 weeks of high-fat diet and L-NAME (HFD+L-NAME) or standard chow (CON) feeding in female mice. **(B)** Exercise capacity as assessed by graded treadmill running, recording total distance travelled (meters) until exhaustion. **(C)** Non-invasive systolic blood pressure (mm Hg) measured via tail-cuff at 6 weeks of HFD-L-NAME treatment. **(G)** Circulating glucose levels (mg/dL) during glucose tolerance testing of *Nnt*^+/+^ and *Nnt*^-/-^ mice.* **(H)** Area-under-curve analysis of glucose tolerance. Statistical significance was assigned via 1-way ANOVA with Tukey’s multiple comparisons test for all pairwise analysis, reporting group mean ± S.E.M. * P < 0.05, ** P < 0.01, *** P < 0.001.

**Supplemental Figure 2.**
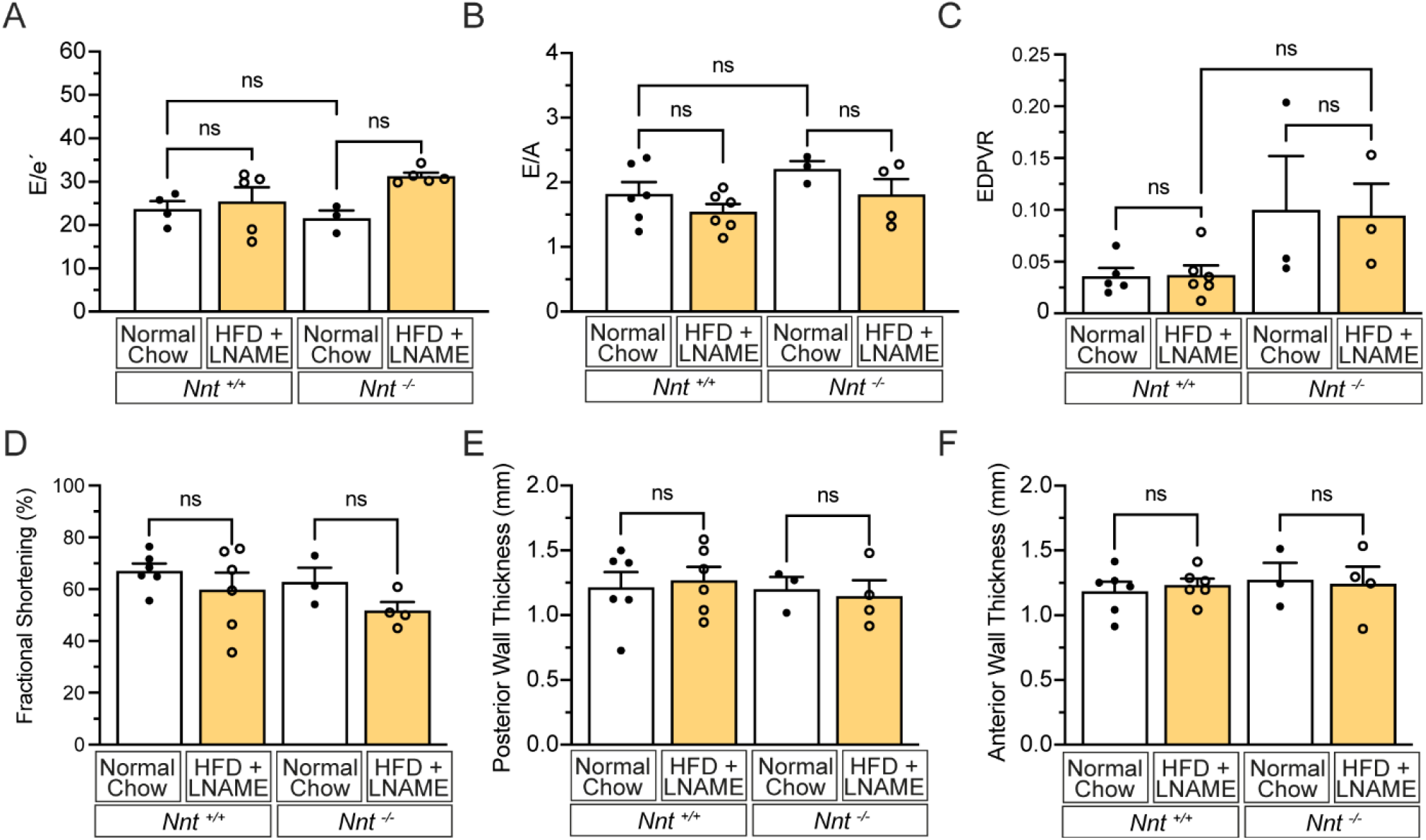
Absence of cardiac phenotype among female mice independent of genotype. **(A)** Schema illustrating the experimental groups used in the study.**(B)** Experimental timeline with concordant endpoints used. **(C)** Body weight (grams) measured after 9 weeks of high-fat diet and L-NAME (HFD+L-NAME) or standard chow (CON) feeding in male mice. **(D)** Exercise capacity as assessed by graded treadmill running, recording total distance travelled (meters) until exhaustion. **(E)** Non-invasive systolic blood pressure (mm Hg) measured via tail-cuff at 6 weeks of HFD+L-NAME treatment. **(F)** Circulating glucose levels (mg/dL) measured via POC glucometer following 4-hr fasting. **(G)** Circulating glucose levels (mg/dL) during glucose tolerance testing of *Nnt*^+/+^ and *Nnt*^-/-^ mice.* **(H)** Area-under-curve analysis of glucose tolerance. Statistical significance was assigned via 1-way ANOVA with Tukey’s multiple comparisons test for all pairwise analysis, reporting group mean ± S.E.M. * P < 0.05, ** P < 0.01, *** P < 0.001.

**Supplemental Figure 3.**
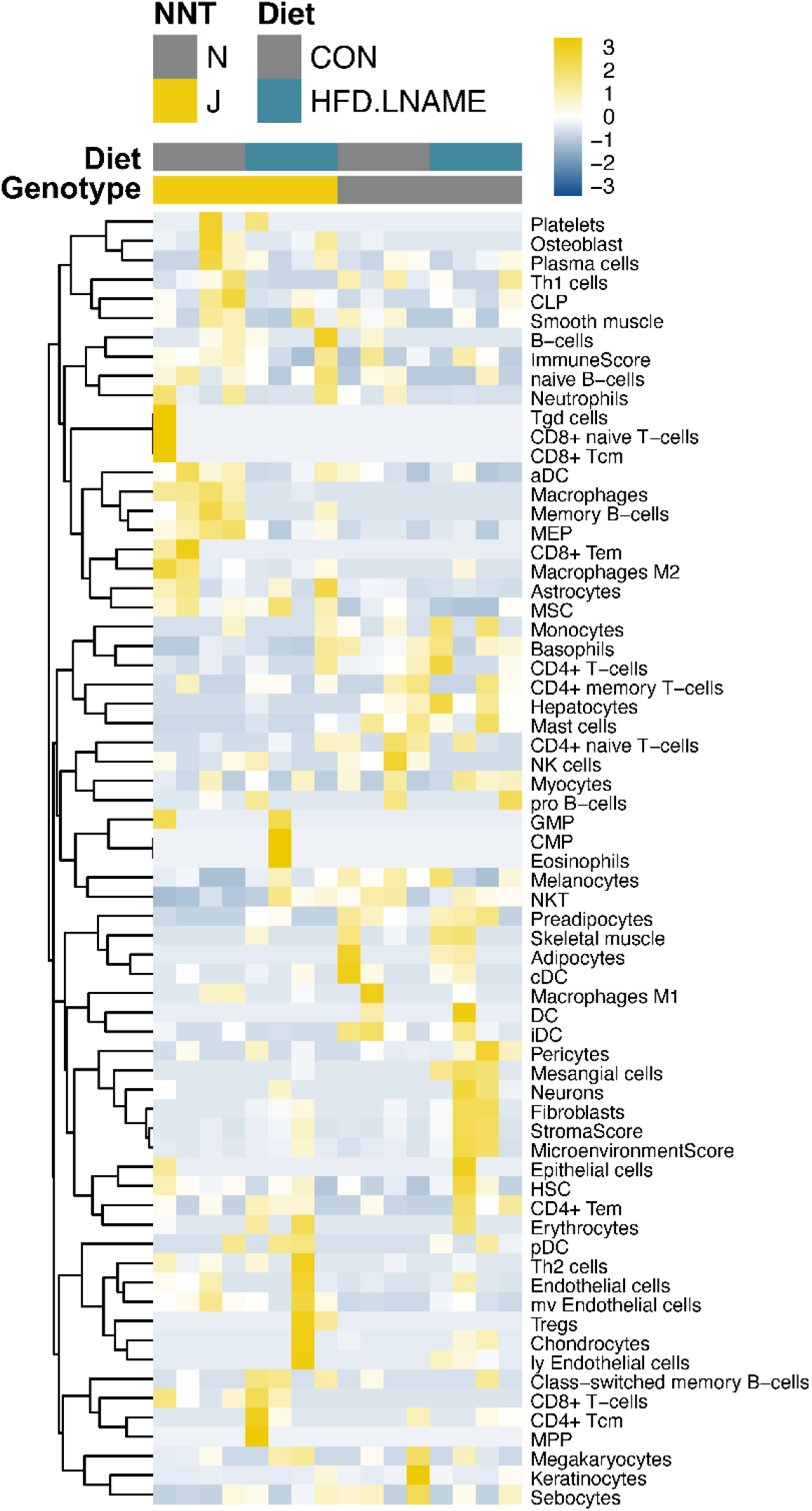
Transcriptomic *In Silico* Immuno/Cellular Deconvolution. *Excell* was used to deconvolute normalized RNA-sequencing count data from left ventricular tissue obtained from mice following 9 weeks of high-fat diet and L-NAME (HFD+L-NAME) or standard chow (CON) feeding.

